# Static and Dynamic Cross-Network Functional Connectivity Shows Elevated Entropy in Schizophrenia Patients

**DOI:** 10.1101/2024.06.15.599084

**Authors:** Natalia Maksymchuk, Juan R. Bustillo, Daniel H. Mathalon, Adrian Preda, Robyn L. Miller, Vince D. Calhoun

**Affiliations:** Tri-Institutional Center for Translational Research in Neuroimaging and Data Science (TReNDS): Georgia State University, Georgia Institute of Technology and Emory University, Atlanta, GA, USA; Department of Psychiatry and Behavioral Sciences, University of New Mexico, Albuquerque, NM, USA; Department of Psychiatry and Behavioral Sciences, Weill Institute for Neurosciences, University of California, San Francisco, CA, USA; Mental Health Service, San Francisco Veterans Affairs Healthcare System, San Francisco, CA, USA; Department of Psychiatry and Human Behavior, University of California, Irvine, CA, USA

**Keywords:** schizophrenia, entropy, brain states, static functional connectivity, dynamic functional connectivity, functional connectivity patterns, mental health, biomarkers, fMRI, image data analysis

## Abstract

Schizophrenia (SZ) patients exhibit abnormal static and dynamic functional connectivity across various brain domains. We present a novel approach based on static and dynamic inter-network connectivity entropy (ICE), which represents the entropy of a given network’s connectivity to all the other brain networks. This novel approach enables the investigation of how connectivity strength is heterogeneously distributed across available targets in both SZ patients and healthy controls. We analyzed fMRI data from 151 schizophrenia patients and demographically matched 160 healthy controls. Our assessment encompassed both static and dynamic ICE, revealing significant differences in the heterogeneity of connectivity levels across available brain networks between SZ patients and healthy controls (HC). These networks are associated with subcortical (SC), auditory (AUD), sensorimotor (SM), visual (VIS), cognitive control (CC), default mode network (DMN) and cerebellar (CB) functional brain domains. Elevated ICE observed in individuals with SZ suggests that patients exhibit significantly higher randomness in the distribution of time-varying connectivity strength across functional regions from each source network, compared to healthy control group. C-means fuzzy clustering analysis of functional ICE correlation matrices revealed that SZ patients exhibit significantly higher occupancy weights in clusters with weak, low-scale functional entropy correlation, while the control group shows greater occupancy weights in clusters with strong, large-scale functional entropy correlation. k-means clustering analysis on time-indexed ICE vectors revealed that cluster with highest ICE have higher occupancy rates in SZ patients whereas clusters characterized by lowest ICE have larger occupancy rates for control group. Furthermore, our dynamic ICE approach revealed that it appears healthy for a brain to primarily circulate through complex, less structured connectivity patterns, with occasional transitions into more focused patterns. However, individuals with SZ seem to struggle with transiently attaining these more focused and structured connectivity patterns. Proposed ICE measure presents a novel framework for gaining deeper insights into understanding mechanisms of healthy and disease brain states and a substantial step forward in the developing advanced methods of diagnostics of mental health conditions.

## 1 Introduction

The advancement of tools designed to provide quantitative biomarkers for various psychiatric disorders is of increasing interest. These tools seek to enhance the diagnosis and screening of the condition, while also offering further insights into the underlying neural mechanisms of mental disorders (Racz et al., 2020). Evaluating properties of brain network connectivity obtained from resting-state (task-free) functional magnetic resonance imaging (rs-fMRI) is widely used for identifying characteristic and reproducible brain activation patterns associated with distinct cognitive and clinical conditions (Allen et al., 2014; Arbabshirani et al., 2013; Damaraju et al., 2014; Du et al., 2020; Li et al., 2020; Liu et al., 2008; Lurie et al., 2020; Miller, Vergara, et al., 2016; Sakoğlu et al., 2010). In contrast to task-based fMRI, rs-fMRI is obtained without external stimuli or tasks, allowing for the capture of the brain’s spontaneous activity during rest. Thus, rs-fMRI allows to explore spatiotemporal organization of the brain on macro-scale level. The primary signal utilized in rs-fMRI is the blood oxygenation-level dependent (BOLD) signal, reflecting alterations in oxygenation levels that are associated with neural activity across various brain regions. From a clinical perspective rs-fMRI provides several advantages. It is a non-invasive technique that is relatively straightforward to administer, placing fewer demands on patients compared to other imaging methods or task-based fMRI paradigms (Alaçam et al., 2023; Arbabshirani et al., 2013; Duda et al., 2023; Iraji et al., 2022; Iraji et al., 2023; Lee et al., 2013), it show robustness in clinical applications even at short scan time (2-5 min) (Duda et al., 2023), as well as it allows to identify individual’s unique functional brain connectivity profile (Finn et al., 2015). This is particularly crucial for clinical populations who may struggle to perform standardized tasks within the scanner.

The traditional approach to functional brain connectivity has involved assuming a static connectivity pattern throughout the data acquisition period (Hutchison, Womelsdorf, Allen, et al., 2013). However, it has been shown that spontaneous BOLD signals recorded during periods of rest display inherent spatiotemporal dynamic organization (Chang & Glover, 2010; Hutchison, Womelsdorf, Gati, et al., 2013; Sakoğlu et al., 2010). Dynamic functional network connectivity (dFNC) is one of the strategies proposed to characterize time-varying brain properties (Sakoğlu et al., 2010). Within this framework, the brain is partitioned into independent networks using a method known as group independent component analysis (ICA) each with its unique temporal profile (Calhoun & Adalı, 2012; Calhoun et al., 2014). The subsequent examination of time-varying changes among component time courses, known as functional network connectivity (FNC), involves calculating cross-correlations between brain networks (components) over time (Calhoun et al., 2014; Jafri et al., 2008). The correlation patterns evolve over time, reflecting fluctuations in neural activity at the macroscopic level and provide insights into how brain networks evolve and interact over different time scales. Afterward, clustering analysis is executed on the time series of correlation patterns to identify matrices representing connectivity “states”. These states are considered to be fundamental to cognition and behavior and useful for characterizing distinct clinical conditions (Calhoun et al., 2014; Hutchison, Womelsdorf, Allen, et al., 2013). Although patterns of both static (calculated over an entire scan) and functional connectivity exhibit sensitivity to individual variations in health and disease, dynamic functional network connectivity provides additional results and is considered to be a more sensitive biomarker when compared to static FNC (Damaraju et al., 2014; Jin et al., 2017; Sakoğlu et al., 2010). Altered dFNC patterns have been observed in an expanding range of neurological and psychiatric disorders compared to control groups (Alaçam et al., 2023; Allen et al., 2014; Damaraju et al., 2014; de Lacy & Calhoun, 2019; de Lacy et al., 2017; Duda et al., 2023; Jin et al., 2017; Lurie et al., 2020; Miller, Vergara, et al., 2016; Sakoğlu et al., 2010).

Schizophrenia (SZ), a prevalent mental disorder affecting around 1% of the world’s population, encompasses a complex array of symptoms that impact cognition, perception, and emotional regulation, often resulting in disruptions to daily functioning (Bhugra, 2005; Wyatt et al., 1995). Ongoing research endeavors aim to elucidate its intricate mechanisms, with a particular focus on comprehending changes in dFNC, which offer invaluable insights into the dynamic brain processes associated with SZ. SZ is characterized by dysconnectivity, which refers to the abnormal functional integration of brain processes. This dysconnectivity implies disrupted communication between different brain regions. Individuals diagnosed with schizophrenia, particularly those exhibiting heightened hallucinatory propensities, exhibit a notable decrease in the dynamic activity of time-varying whole-brain network connectivity patterns (Miller, Vergara, et al., 2016; Miller, Yaesoubi, et al., 2016). Also, SZ patients showed a reduction in temporal autocorrelations, reduced multifractality and increased self-similarity (Alamian et al., 2022). Furthermore, SZ affects the sensitivity of intra-network connectivity to broader functional brain interactions (Miller, Vergara, et al., 2016). In healthy subjects, patterns of connectivity within the intra-auditory-visual-sensorimotor networks (AVSN) show responsiveness to variations in network relationships across various domains. Conversely, individuals with SZ exhibit isolated intra-AVSN connectivity, which does not influence or respond to changes in network relationships within domain pairs containing at least one non-AVSN functional domain (Miller, Vergara, et al., 2016). The neural mechanisms of dysconnectivity observed in SZ patients remain to be fully unraveled, and research continues to investigate their dynamics and clinical significance. Schizophrenia presents as a complex disorder exhibiting disrupted brain network interactions at both static and dynamic levels, thus requiring sophisticated approaches to reveal its underlying neural mechanisms.

In recent years, there has been a notable increase in empirical studies with a focus on integration of both structural and functional connectivity analyses with information theory offering a powerful framework for advancing our understanding of brain organization (Poza et al., 2021). Metrics originating from information theory, particularly those linked with entropy, have shown their ability in extracting meaningful information from underlying brain networks, in both healthy and mental disorder state (Poza et al., 2021). Thus, current study (Blair et al., 2024) tracked subject trajectories in dynamic functional connectivity state space during brain scans evaluating entropy production along each dimension of the proposed basis space. Authors found that schizophrenia patients demonstrate lower entropy, suggesting simpler trajectories compared to healthy controls.

In present work we introduced novel measure combining FNC and information theory approaches – inter-network connectivity entropy (ICE), entropy of distribution of time-varying connectivity strength across functional brain regions. We investigated static and dynamic ICE across 53 functional intrinsic brain networks extracted from rs-fMRI data from 311 subjects, including 151 schizophrenia patients and 160 healthy controls, to discern potential differences in ICE between SZ patients and controls and to determine functional brain networks that exhibit those differences and evaluate whether they manifest as higher or lower values in SZ patients relative to controls. Higher values of ICE indicate higher randomness and more heterogeneity of connectivity levels across available networks whereas lower ICE values are evidence of less randomness and more concentration (less heterogeneity) in connectivity levels. In addition, we performed C-means fuzzy clustering on functional ICE correlation matrices to uncover potential differences in functional entropy correlation between and within intrinsic brain networks in both the SZ patient and control groups. Furthermore, we employed k-means clustering of time-indexed ICE vectors to identify characteristic ICE states and their occupancies for each group. Our approach provides new insights into unraveling the neural mechanisms of dysconnectivity in SZ patients and for developing advanced biomarkers of the of mental health conditions.

## 2 Materials and Methods

### 2.1 fMRI Data

We used resting-state fMRI data collected from a total of 311 participants, comprising 160 healthy controls (HC) and 151 individuals diagnosed with SZ, matched for age and gender. The data were acquired as part of the multi-site fBIRN project (Potkin & Ford, 2009). Participants were directed to keep their eyes closed throughout the scans. Data collection occurred every 2 seconds (TR) for a total of 160 TRs, equivalent to 5.33 minutes. The data underwent preprocessing using a standard pipeline, as detailed in (Damaraju et al., 2014; Du et al., 2020), and underwent decomposition with group-independent component analysis. This process yielded 100 group-level functional network spatial maps along with their corresponding timecourses (**Figure 1**). Among these components, 53 were identified as intrinsic connectivity networks (ICNs), in accordance with the methods described in earlier publications (Damaraju et al., 2014; Du et al., 2020). Subject-specific spatial maps and temporal profiles were obtained using spatiotemporal regression. The temporal profiles of each subject’s ICNs were detrended, orthogonally aligned with motion parameters, and despiked. Detailed description of data collection, estimation of the functional networks, their functional connectivity and number of temporally independent sources are provided in (Blair et al., 2024; Du et al., 2020).

**Figure 1.**
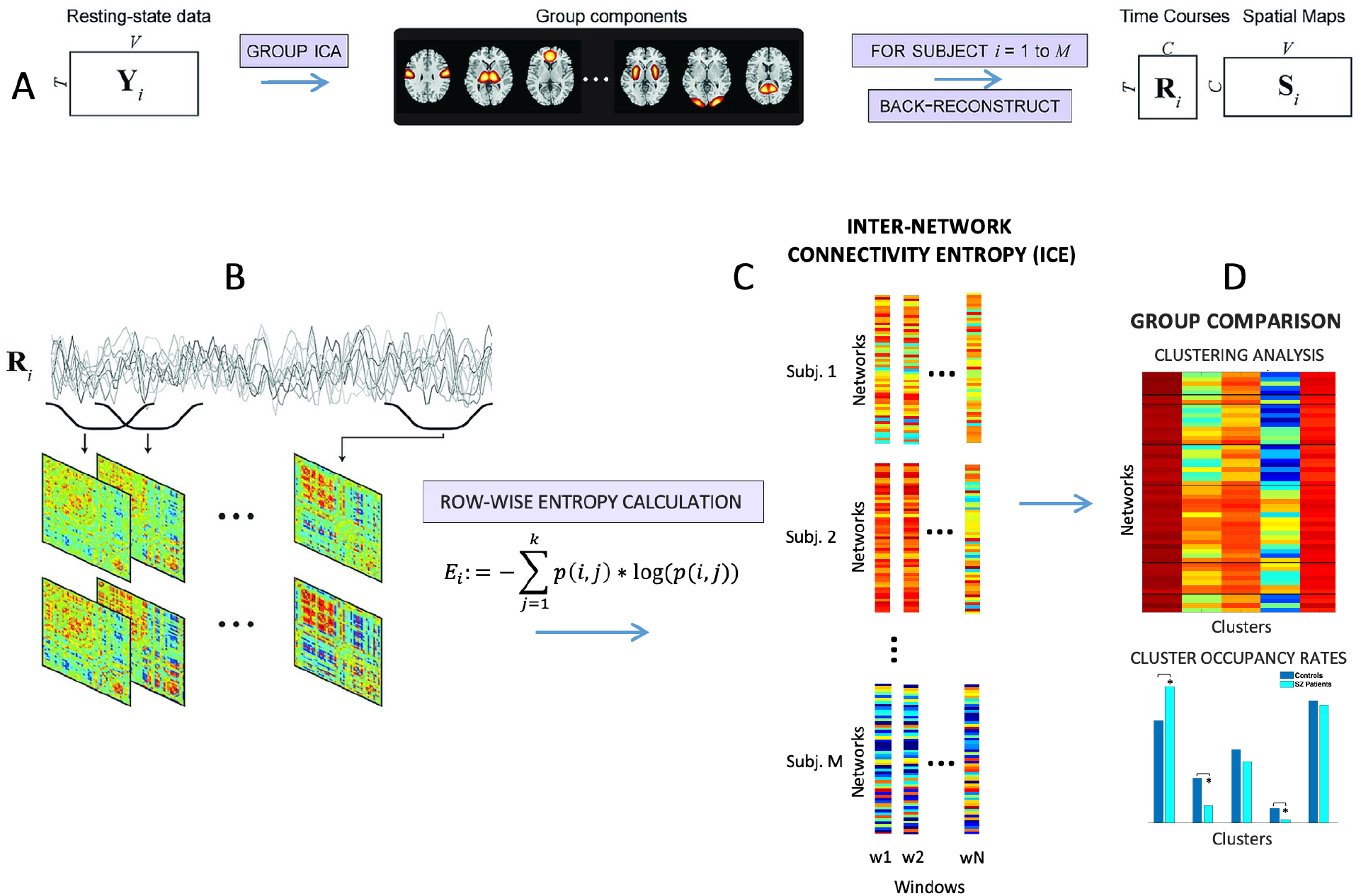
Schematics of main steps of analysis, modified from R. L. Miller *et al*., “Higher Dimensional Meta-State Analysis Reveals Reduced Resting fMRI Connectivity Dynamism in Schizophrenia Patients,” *PLoS One*, 2016. (**A**) Decomposition of resting-state fMRI data with GICA into network spatial maps and corresponding time courses. (**B**) Obtaining dynamic functional connectivity matrices for each of subject. (**C**) Computing intra-network connectivity entropy (ICE) for controls and SZ patients from dynamic functional connectivity matrices. After we applied clustering algorithms and regression analysis to determine group difference between SZ and HC (**D**).

### 2.2 Inter-network Connectivity Entropy (ICE)

First, we calculated network connectivity distributions from dFNC matrices. Next, we determined the entropies of these distributions (inter-network connectivity entropies (ICE)). Network connectivity distributions and ICE were calculated in static and dynamic ways, obtaining ICE aggregated over all time windows, static ICE (SICE), and window-wise ICE or dynamic ICE (DICE).

For each network *i*, we look at its connectivities *c*(*i, j*)obtained from dFNC matrices across *j* ≠ *i* as a distribution of connectivity strengths across functional intrinsic brain networks. We map each *c*(*i, j*)to a non-negative translate denoted as *c′* (*i, j*)= *c*(*i, j*)– *C*_*min*_, where *C*_*min*_ minimal connectivity on a global level. A probability distribution for each network *i* was computed as *P*_*i*_ consisting of the sequence {*p*(*i*, 1), *p*(*i*, 2), … *p*(*i, k*)}, *j* ≠ *i*, where 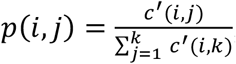, is a summed connectivity of component to each other network rescaled to be a distribution, *k* is a number of components (networks). After that we computed the connectivity entropy these distributions for every network *i* a*s* 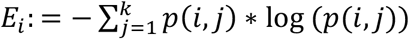.

We obtained tensors of dynamic and static ICE values in dimensions of 53×137x311 and 53×311, respectively. Here, 53 represents the number of functional intrinsic networks, 137 indicates the number of time windows, and 311 signifies the number of subjects. Functional entropy correlation matrices of dimensions 53×53 for both SZ patients and controls were generated by autocorrelation of the 53×137 matrices of DICE for each subject. All computations and data analyses were performed utilizing custom MATLAB scripts. Connectograms depicting functional ICE correlations were generated using GIFT toolbox function ‘icatb_plot_connectogram’ (http://trendscenter.org/software/gift) (Iraji et al., 2021) and Neuromark fMRI 1.0 template (Du et al., 2020).

### 2.3 Clustering Analysis

The C-means fuzzy clustering was performed on functional ICE correlation matrices of all subjects with the Euclidean distance, 500 iterates, fuzziness parameter equal 1.05. The set of functional ICE correlation matrices was segmented into five clusters, with their centroids serving as basis correlation patterns. Cluster occupancy weights were derived from the fuzzy partition matrix, which contains the percentage of cluster membership for each observation.

The k-means clustering algorithm was applied to the time-indexed entropy vectors partitioning data into five different clusters using Euclidean distance, 500 iterates, and 50 replicates followed by assessment of subject-level cluster occupancy rates and dwell time for both SZ patients and HC. Number of clusters was established using the elbow criterion. Both k-means and c-means clustering utilized MATLAB’s functions.

### 2.4 Statistics

A linear regression model and two-sample t-test were employed to assess the impact of schizophrenia on ICE. The reported p-values underwent correction for multiple comparisons using FDR (false discovery rate) at *α*_*FDR*_ = 0.05. The regression model accounts for potential confounding variables such as age, gender, and mean frame displacement (motion). The diagnosis variable is binary, where ‘1’ represents SZ and ‘0’ represents HC. Therefore, a positive regression coefficient for diagnosis indicates a positive correlation with SZ, while a negative value of regression coefficient for diagnosis indicates a negative correlation with SZ.

## 3 Results

### 3.1 SZ patients tend to display higher static and dynamic ICE across the majority of intrinsic brain connectivity networks when contrasted with healthy controls

In our study, our goal was to examine heterogeneity in connectivity strength distributions across intrinsic connectivity brain networks in both SZ patients and healthy controls. To accomplish this, we computed the entropy of connectivity strength distributions within functional brain regions of each source network, termed as intra-network connectivity entropy (ICE). Among the 53 functional brain networks examined, 36 exhibited statistically significant difference in static ICE between SZ patients and controls (p≤0.0274 (FDR)) (**Table 1**). These implicated networks encompass diverse functional brain domains, such as with subcortical (SC), auditory (AUD), visual (VIS), sensorimotor (SM), cognitive control (CC), default mode networks (DMN) and cerebellar (CB). Furthermore, dynamic ICE showed significant differences in 41 out of the 53 functional networks (p≤0.0379 (FDR)) affecting same functional brain domains (**Table 1**). Mean static and mean dynamic ICEs computed across 53 functional connectivity networks for both healthy controls and SZ patients are illustrated in **Figures 2 and 3**.

**Table 1.**
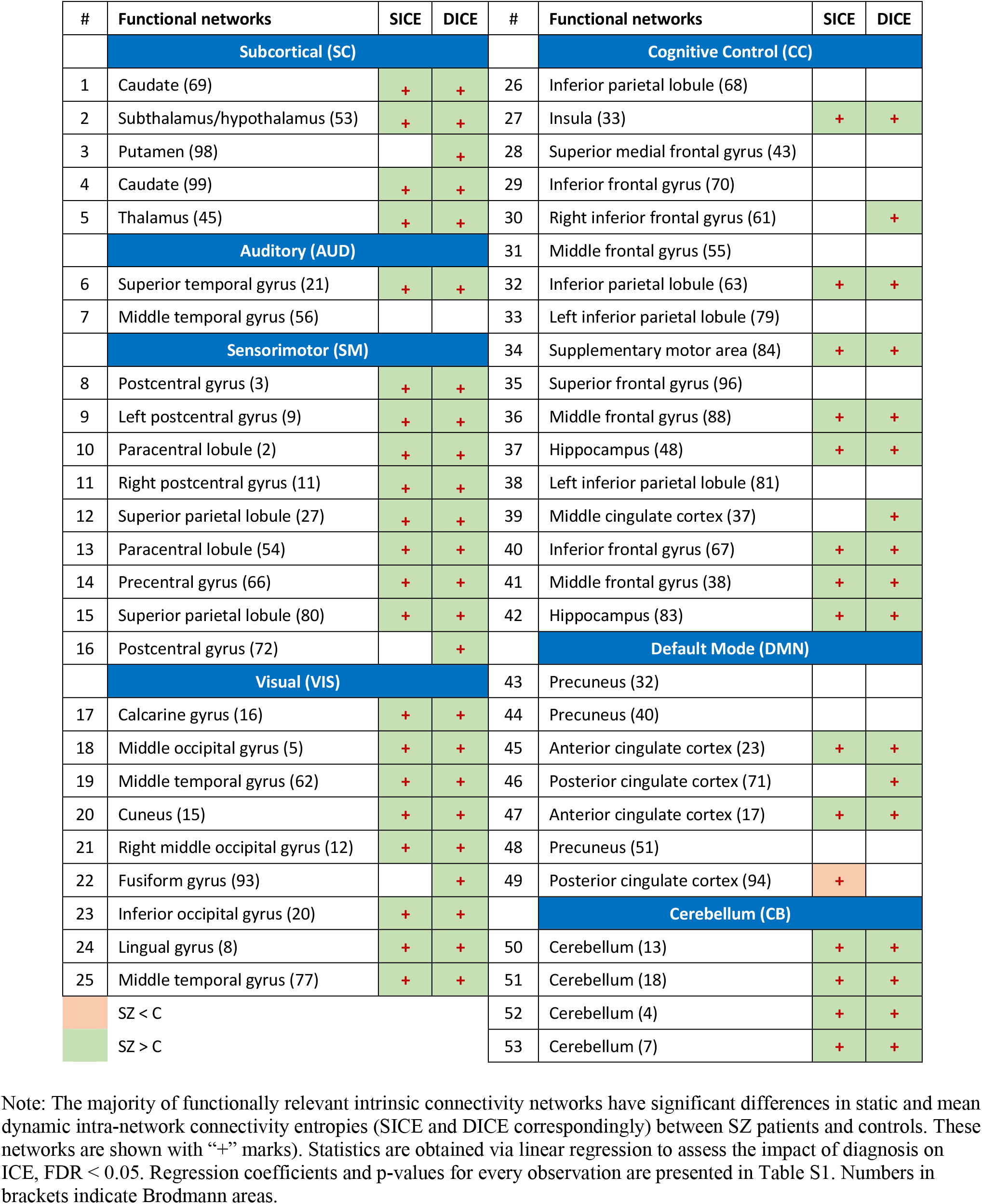
Mean ICE associated with intrinsic connectivity networks.

| # | Functional networks | SICE | DICE | # | Functional networks | SICE | DICE |
| --- | --- | --- | --- | --- | --- | --- | --- |
|  | **Subcortical (SC)** |  |  |  | **Cognitive Control (CC)** |  |  |
| 1 | Caudate (69) | + | + | 26 | Inferior parietal lobule (68) |  |  |
| 2 | Subthalamus/hypothalamus (53) | + | + | 27 | Insula (33) | + | + |
| 3 | Putamen (98) |  | + | 28 | Superior medial frontal gyrus (43) |  |  |
| 4 | Caudate (99) | + | + | 29 | Inferior frontal gyrus (70) |  |  |
| 5 | Thalamus (45) | + | + | 30 | Right inferior frontal gyrus (61) |  | + |
|  | **Auditory (AUD)** |  |  | 31 | Middle frontal gyrus (55) |  |  |
| 6 | Superior temporal gyrus (21) | + | + | 32 | Inferior parietal lobule (63) | + | + |
| 7 | Middle temporal gyrus (56) |  |  | 33 | Left inferior parietal lobule (79) |  |  |
|  | **Sensorimotor (SM)** |  |  | 34 | Supplementary motor area (84) | + | + |
| 8 | Postcentral gyrus (3) | + | + | 35 | Superior frontal gyrus (96) |  |  |
| 9 | Left postcentral gyrus (9) | + | + | 36 | Middle frontal gyrus (88) | + | + |
| 10 | Paracentral lobule (2) | + | + | 37 | Hippocampus (48) | + | + |
| 11 | Right postcentral gyrus (11) | + | + | 38 | Left inferior parietal lobule (81) |  |  |
| 12 | Superior parietal lobule (27) | + | + | 39 | Middle cingulate cortex (37) |  | + |
| 13 | Paracentral lobule (54) | + | + | 40 | Inferior frontal gyrus (67) | + | + |
| 14 | Precentral gyrus (66) | + | + | 41 | Middle frontal gyrus (38) | + | + |
| 15 | Superior parietal lobule (80) | + | + | 42 | Hippocampus (83) | + | + |
| 16 | Postcentral gyrus (72) |  | + |  | **Default Mode (DMN)** |  |  |
|  | **Visual (VIS)** |  |  | 43 | Precuneus (32) |  |  |
| 17 | Calcarine gyrus (16) | + | + | 44 | Precuneus (40) |  |  |
| 18 | Middle occipital gyrus (5) | + | + | 45 | Anterior cingulate cortex (23) | + | + |
| 19 | Middle temporal gyrus (62) | + | + | 46 | Posterior cingulate cortex (71) |  | + |
| 20 | Cuneus (15) | + | + | 47 | Anterior cingulate cortex (17) | + | + |
| 21 | Right middle occipital gyrus (12) | + | + | 48 | Precuneus (51) |  |  |
| 22 | Fusiform gyrus (93) |  | + | 49 | Posterior cingulate cortex (94) | + |  |
| 23 | Inferior occipital gyrus (20) | + | + |  | **Cerebellum (CB)** |  |  |
| 24 | Lingual gyrus (8) | + | + | 50 | Cerebellum (13) | + | + |
| 25 | Middle temporal gyrus (77) | + | + | 51 | Cerebellum (18) | + | + |
|  | SZ < C |  |  | 52 | Cerebellum (4) | + | + |
|  | SZ > C |  |  | 53 | Cerebellum (7) | + | + |

Note: The majority of functionally relevant intrinsic connectivity networks have significant differences in static and mean dynamic intra-network connectivity entropies (SICE and DICE correspondingly) between SZ patients and controls. These networks are shown with “+” marks). Statistics are obtained via linear regression to assess the impact of diagnosis on ICE, FDR < 0.05. Regression coefficients and p-values for every observation are presented in Table S1. Numbers in brackets indicate Brodmann areas.

**Figure 2.**
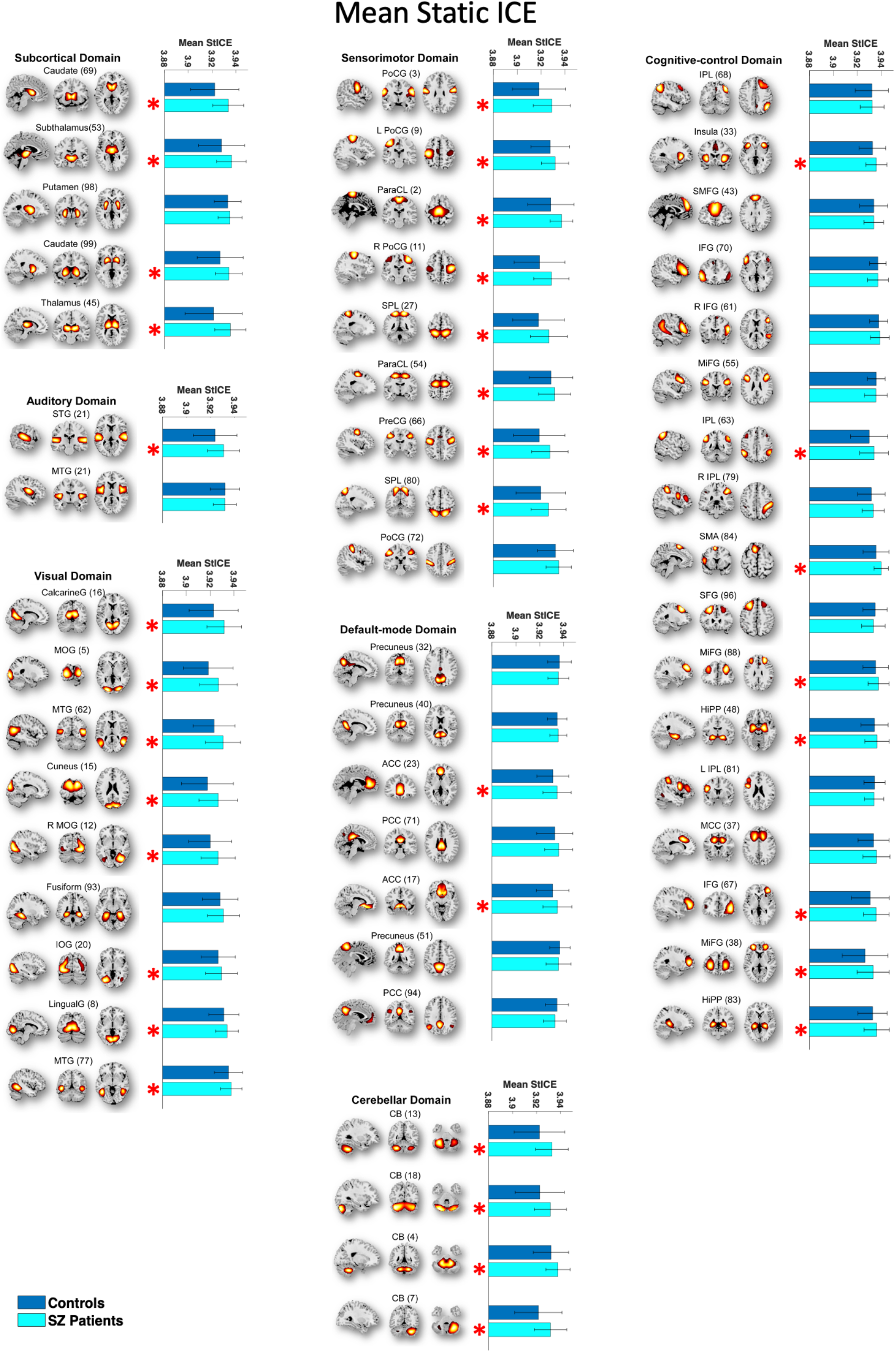
The majority functional brain networks demonstrate significantly higher mean static ICE in SZ patients compared to control group. Networks that have significant differences in mean static intra-network connectivity entropies (SICE) between SZ patients and controls are shown with red “*” marks. The statistical results were acquired from the diagnosis term in univariate multiple regression models.

**Figure 3.**
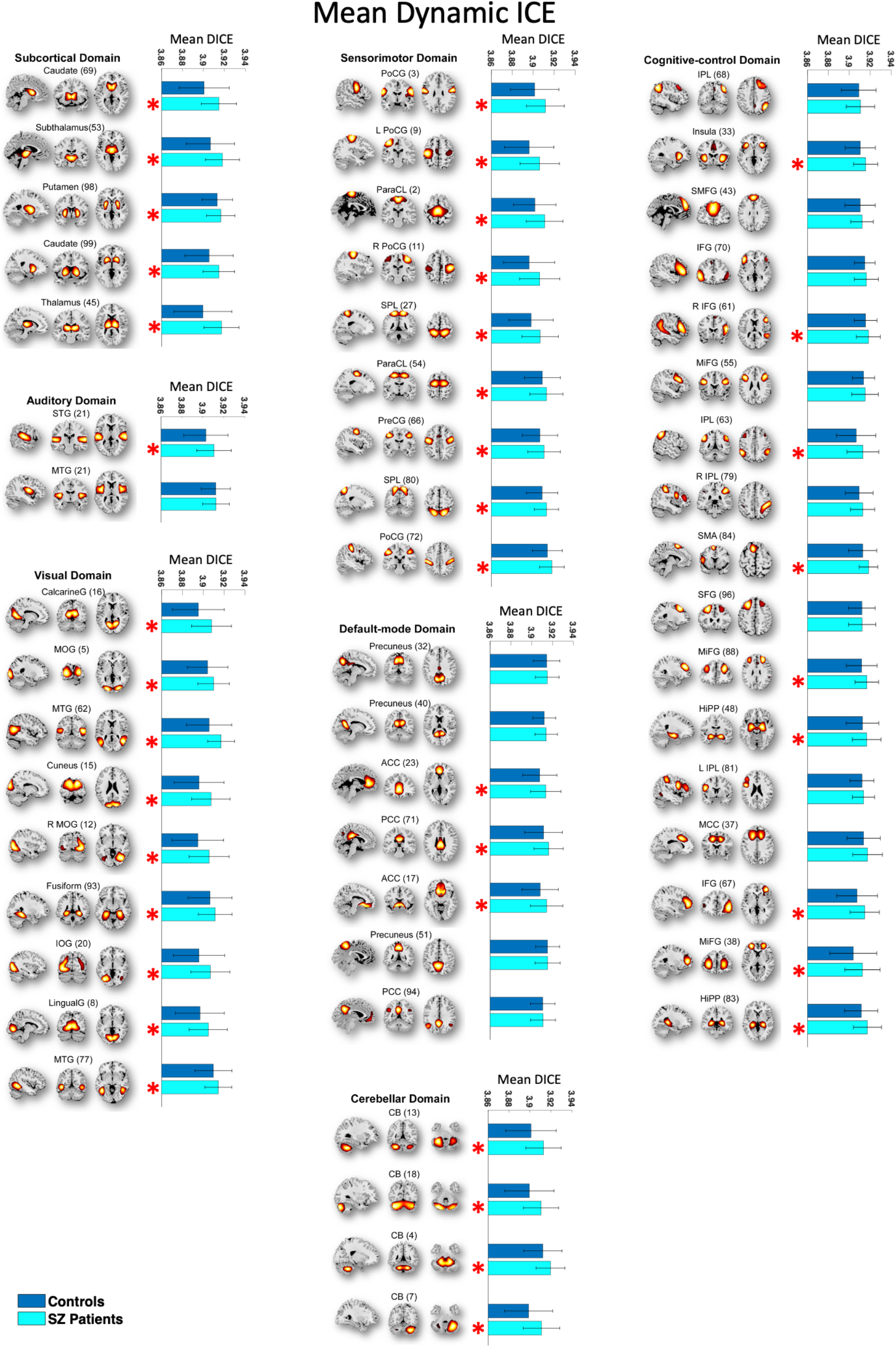
The majority of functional brain networks demonstrate significantly higher mean dynamic ICE (DICE) in SZ patients compared to control group. Networks that have significant differences in mean dynamic intra-network connectivity entropies (DICE) between SZ patients and controls are shown with red “*” marks. The statistical results were acquired from the diagnosis term in univariate multiple regression models.

Next, we assessed whether SZ patients exhibit higher or lower levels of ICE compared to healthy controls. Except for the posterior cingulate cortex, all networks with significant differences in dynamic ICE between patients and controls demonstrated higher ICE in schizophrenia patients compared to controls. While the posterior cingulate cortex network demonstrated higher static ICE in HC, no statistically significant difference in dynamic ICE was observed between SZ patients and HC in this network.

To investigate the effects of age and gender on ICE, we employed a linear regression model while correcting for multiple comparisons. Our analysis indicated that gender does not significantly affect heterogeneity of intra-network connectivity strength distribution for both static and dynamic measures, whereas age has statistically significant effect on Precuneus intrinsic connectivity network for static ICE measure. Additionally, we investigated the effect of the composite cognitive score and the combined effect of the composite cognitive score and diagnosis (composite cognitive score by diagnosis interaction) on ICE group differences. To this end, we added terms for the composite cognitive score and the composite cognitive score by diagnosis interaction to the regression model. The regression analysis showed that there was no statistically significant effect of either the composite cognitive score or the interaction of the composite cognitive score and diagnosis on both static and dynamic ICE.

### 3.2 SZ patients and control group have distinct distribution of ICE across a variety of intrinsic connectivity networks

Mean values of dynamic ICE computed across windows and subjects provide limited information. Therefore, we examined the distributions of dynamic ICE across different subjects for all networks with statistically significant difference between patients and healthy controls. Six representative histograms of dynamic and static ICE for control and SZ groups are shown in **Figures 4 and 5**. The histograms are left-skewed for both patients and controls whereas SZ histograms have bulk of the mass at the higher end in the distributions compared to controls. Among all 41 networks with p≤0.0379 (FDR), the distributions associated with SZ patients were shifted toward higher connectivity entropies compared to controls. This result is consistent and complementary with the findings presented in the previous section, which described a higher mean ICE in SZ patients.

**Figure 4.**
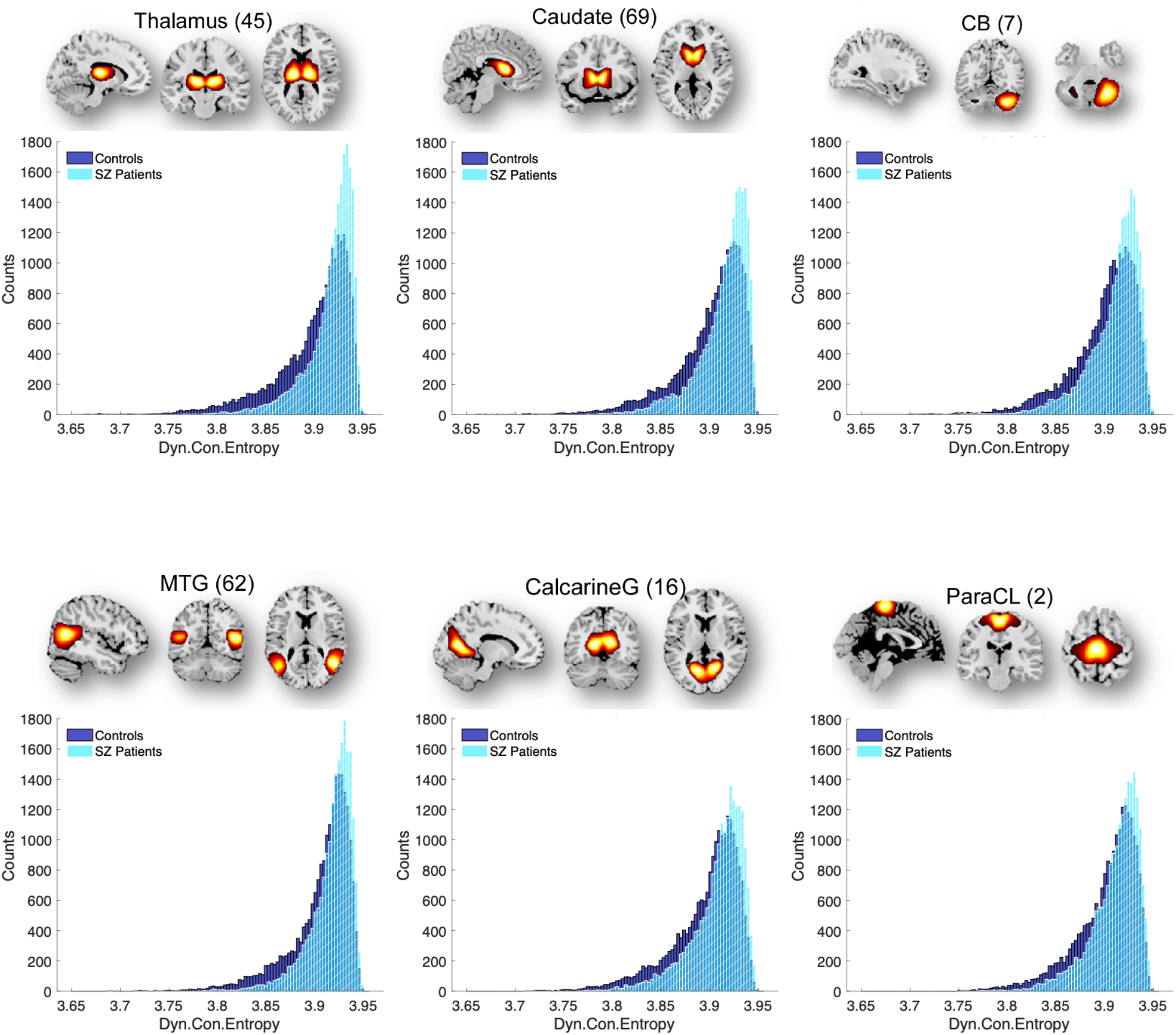
The DICE histograms characterizing SZ patients are skewed towards higher connectivity entropies and contain a larger portion of the mass at the higher end compared to the control group. Six representative functional brain networks Thalamus (SC), Caudate (SC), Cerebellum 4 (CB), Calcarine gyrus (VIS), Middle temporal gyrus (VIS) and Paracentral lobule (SM) with significant difference in dynamic ICE with corresponding p-values: 2.56*10^−10^, 1.27*10^−8^, 2.98*10^−8^, 1.29*10^−7^, 2.28*10^−6^, 4.90*10^−6^. Distributions were obtained for ICE aggregated over all windows and subjects of each group.

**Figure 5.**
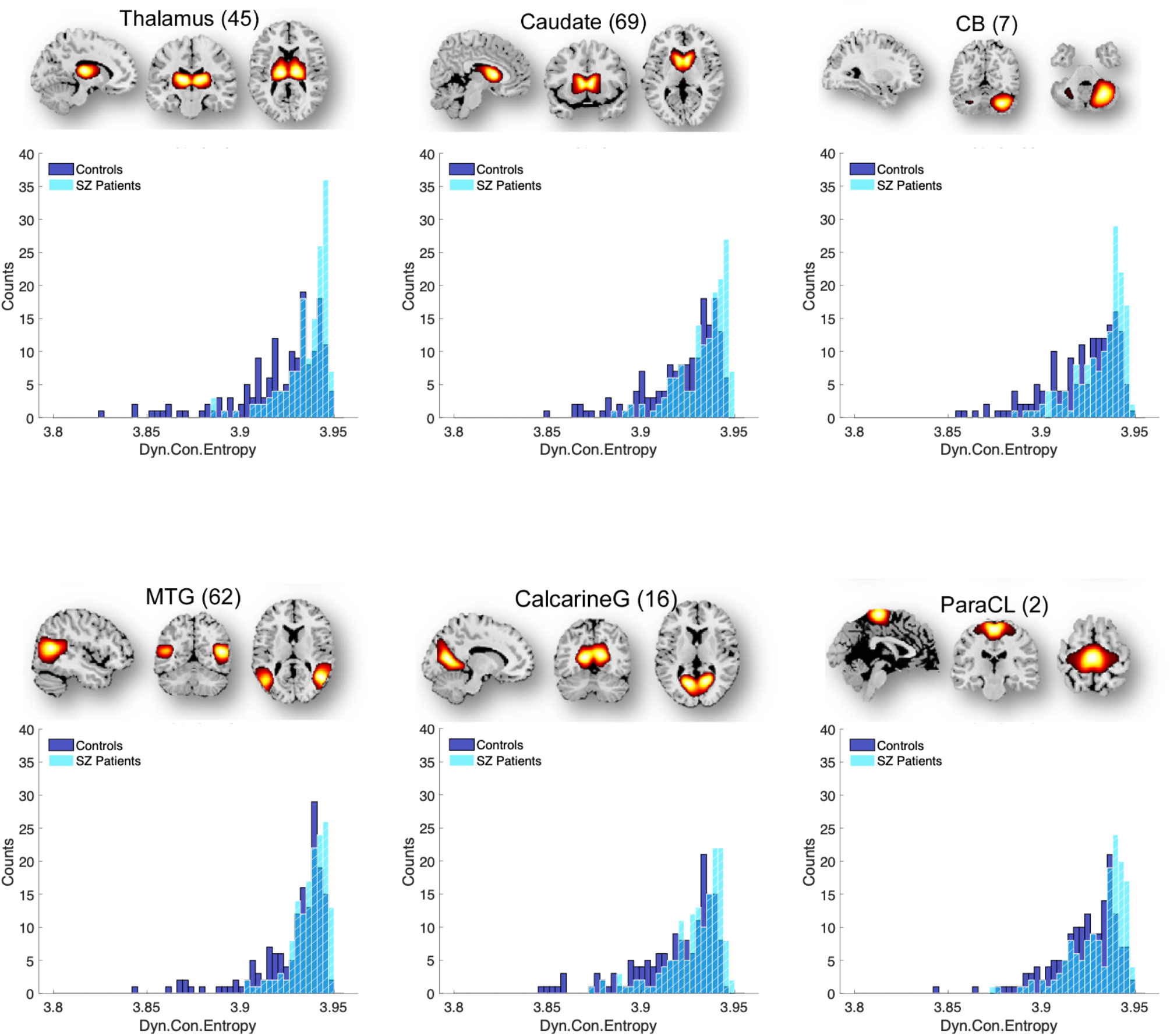
Similarly to DICE histograms, SICE histograms characterizing SZ patients are skewed towards higher connectivity entropies and contain a larger portion of the mass at the higher end compared to the control group. Same as in Figure 4 six representative functional brain networks are Thalamus (SC), Caudate (SC), Cerebellum 4 (CB), Calcarine gyrus (VIS), Middle temporal gyrus (VIS) and Paracentral lobule (SM) with significant difference in dynamic ICE with corresponding p-values: 4.22*10^−9^, 3.96*10^−8^, 6.77*10^−8^, 1.66*10^−6^, 2.61*10^−6^, 8.16*10^−6^. Distributions were obtained averaging ICE over time windows for every subject of each group.

### 3.3 SZ patients have lower variability in intra-network connectivity strength distribution over time compared to the control group

To explore the variability in network connectivity strength distribution over time, we examined the standard deviations (STDs) of the dynamic ICEs across all intrinsic functional brain networks. 46 of 53 functionally relevant intrinsic connectivity networks (shown with red “*” marks in **Figure 6**) have significantly higher STD of the DICE in healthy controls compared to SZ patients. All functional networks with significant differences in SICE and DICE between SZ patients and controls, except posterior cingulate cortex, characterized with high variability in network connectivity strength distribution over time. Moreover, intra-network connectivity in SZ patients exhibits a more uniform distribution, showing relatively consistent temporal patterns, rather than displaying high average entropy driven by specific periods of elevated entropy that skew the average upwards.

**Figure 6.**
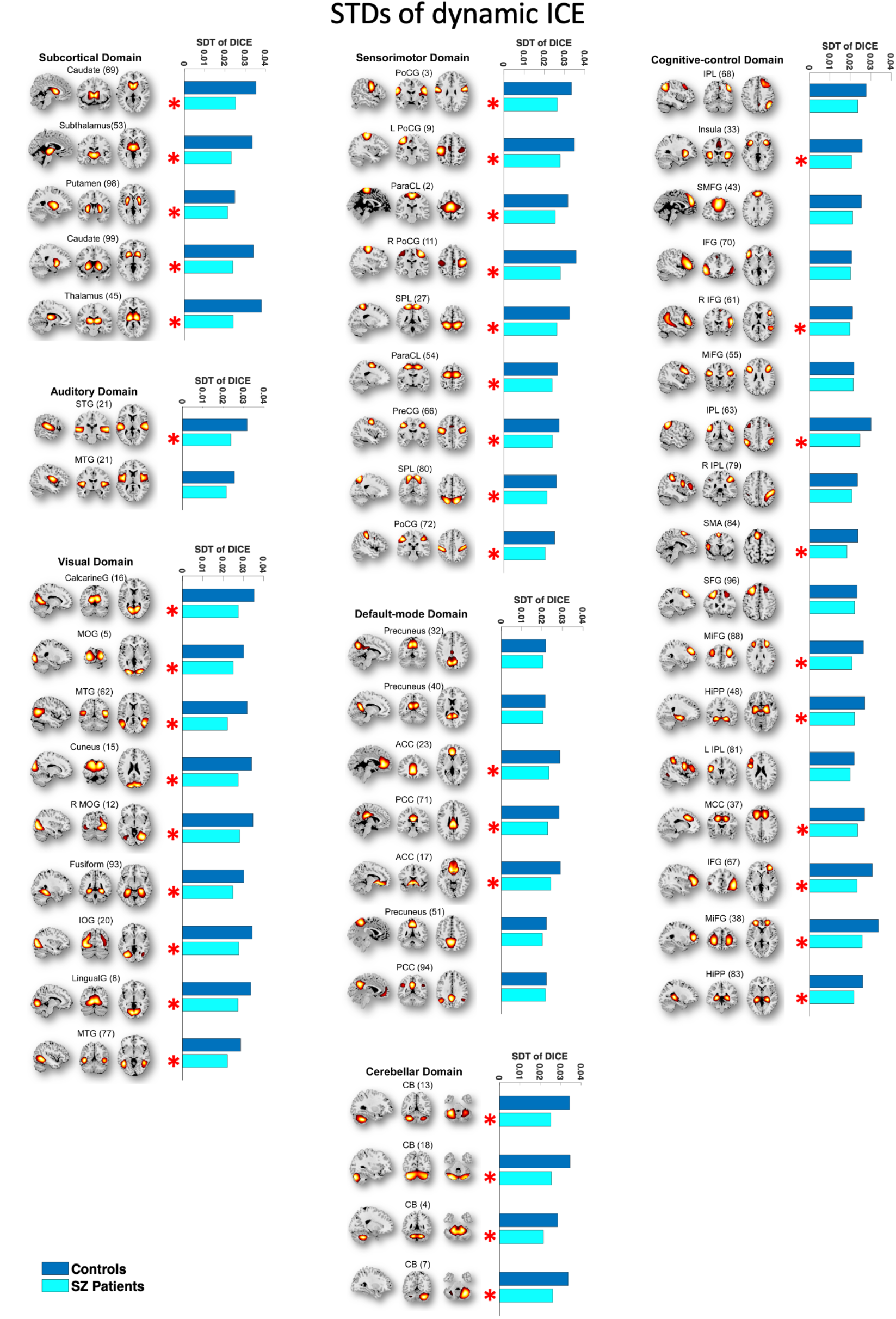
46 out of 53 functionally relevant intrinsic connectivity networks exhibit significantly lower STDs of DICE in SZ patients compared to healthy controls. These networks are shown with “*” marks. Standard deviations were calculated for seven brain domains comprising of 53 functional networks. The majority of these 46 networks have significant differences in SICE and DICE between SZ patients and controls.

### 3.4 SZ patients have distinct ICE patterns in SC, AUD, SM, VIS and CB brain domains compared to healthy controls

To explore correlation of ICE between different brain domains and find potential difference in ICE patterns between SZ patients and control group we investigated whole-brain subject-level functional entropy correlation matrices obtained on time courses of ICE for each component. Averaged functional ICE correlation matrices for both patients and healthy controls and their difference are presented in **Figure 7, A,B** correspondingly. Notably that all network show either positive or no correlation in ICE for both groups. Significant difference is observed in SC, AUD, SM, VIS and CB domains where SZ patients have reduced correlation of ICE between and within networks of these domains, when compared to control group (**Figure 7, C**) what is also depicted on connectograms (**Figure 7, D,E)**.

**Figure 7.**
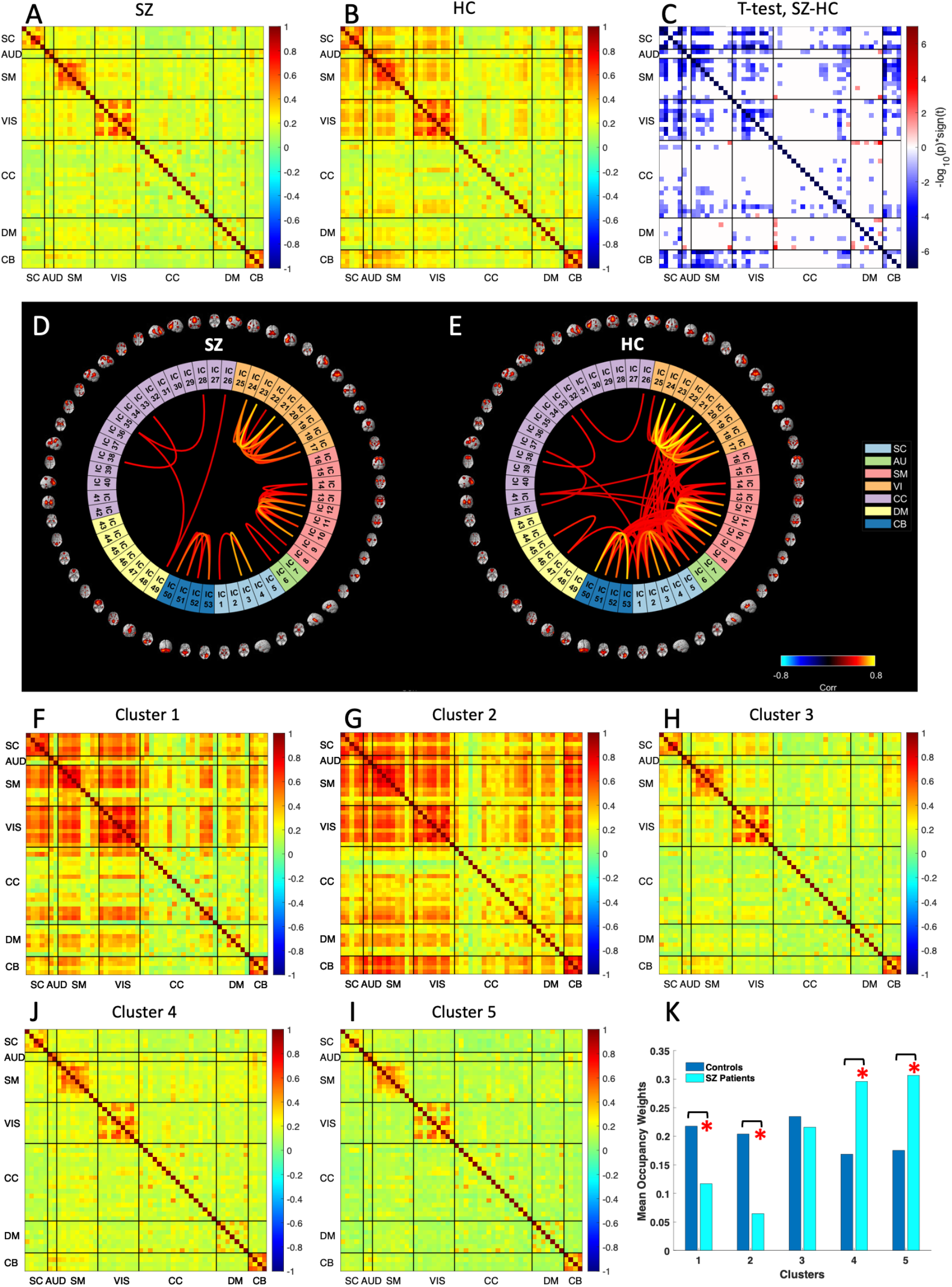
**(A, B)** SZ patients exhibit reduced correlation of ICE between brain regions compared to controls. Mean functional entropy correlation matrices obtained from dynamic ICE for SZ patients and control group correspondingly. **C** The group difference (SZ–HC) in functional entropy correlation matrices. Values are plotted as −log10(p-value)*sign(t-value), where statistics are obtained via t-test across diagnosis groups, FDR < 0.05. The graph displays only the p-values that correspond to statistical significance. **D** and **E** illustrate connectograms derived from mean functional entropy correlation matrices for both the patient and control groups. Connections with correlation values lower than 0.4 are omitted on the connectograms. The c-means algorithm is utilized to cluster functional entropy correlation matrices obtained for all subjects, resulting in the identification of five cluster centroids (**F-I**). **K** Occupancy weights across five clusters for SZ and healthy control group.

Next, we performed clustering analysis of these functional entropy correlation patterns using C-means clustering approach (**Figure 7, F-I**). As a result, we obtained two clusters with strong, large-scale functional entropy correlation, two clusters with weak, low-scale correlation and one cluster with medium functional entropy correlation. Clusters with strong, large-scale functional entropy correlation have larger cluster occupancy weights for controls, whereas clusters with low-scale functional entropy correlation are more occupied by SZ patients (**Figure 7, K**). The results are consistent with FNC clusters for SZ and control groups (Damaraju et al., 2014).

### 3.5 SZ patients and controls exhibit different occupancy rates for clusters exhibiting distinct dynamic ICE patterns

In addition, we performed k-means clustering on the time-indexed ICE vectors. We obtained 5 cluster centroids, that have different ICE patterns (**Figure 8A**). Two of them (1 and 5) are characterized with high entropy and another two (2 and 4) have low entropy values. Noticeably that clusters with high entropy exhibit high ICE across all 53 components, while clusters with low entropy display larger variability in ICE among functional brain networks. Interestingly that SC, AUD, VIS and CB brain domains of clusters 2 and 4 are characterized with lower ICE values compared to other functional brain networks.

**Figure 8.**
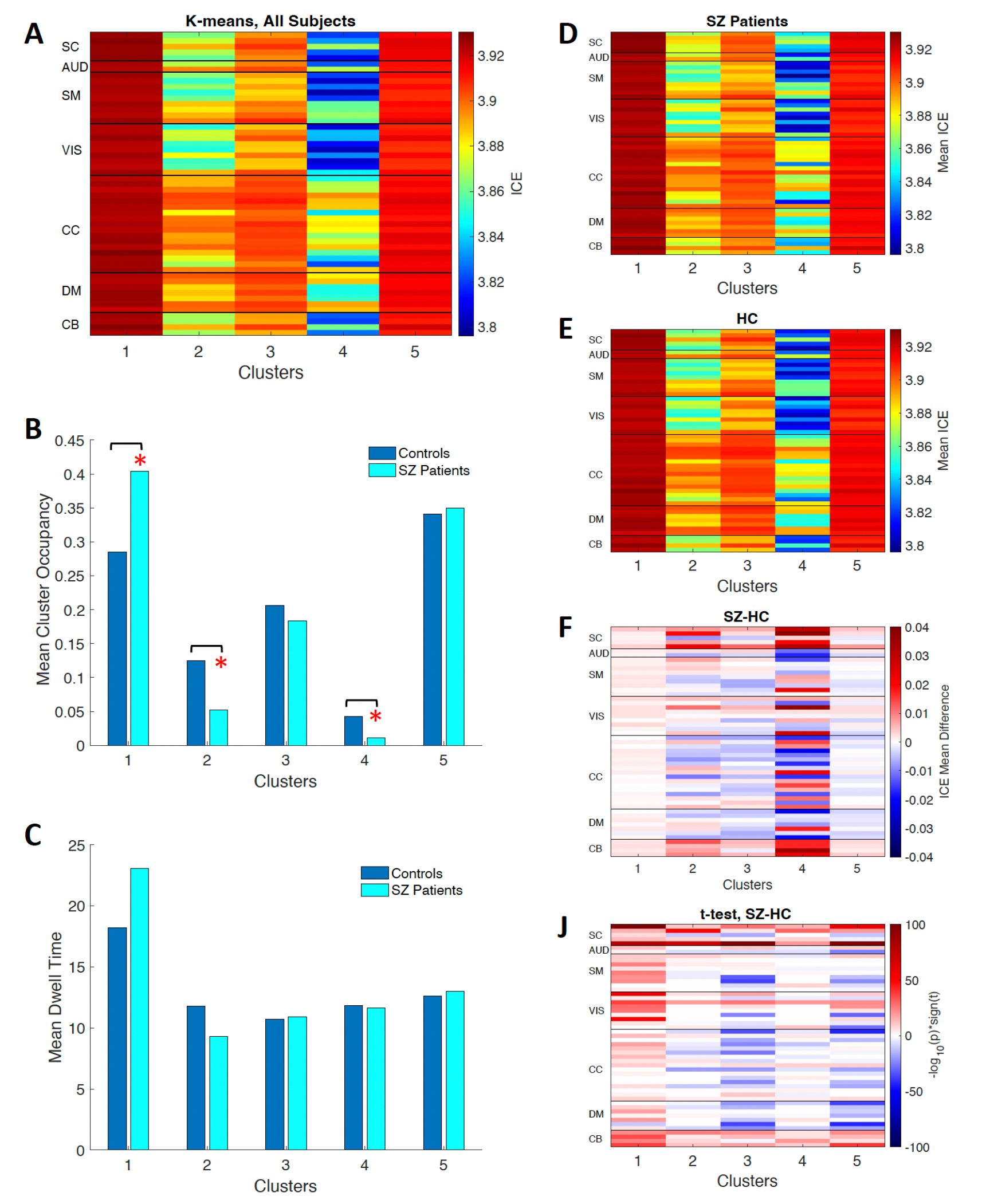
SZ patients are characterized by distinct cluster occupancy rates and different ICE patterns compared to controls (**A**) The k-means clustering algorithm was applied to cluster dynamic ICE obtained for all subjects, resulting in the 5 cluster centroids. (**B**) Mean cluster occupancy rates for all obtained clusters. Cluster 1 with highest entropy across all networks is more occupied by SZs than controls, whereas cluster 4 with lowest entropy is occupied more by healthy controls. (**C**) Mean dwell time associated with each cluster. The mean DICE values corresponding to the five clusters on are shown for healthy controls (**D**) and SZ patients (**E**) and their difference (**F**). (**J**) The group difference (SZ–HC) in mean DICE values for each ICN of each cluster. The values are plotted as −log10(p-value)*sign(t-value), where statistics are obtained via t-test across diagnosis groups, FDR < 0.05. The graph displays only the p-values that correspond to statistical significance.

Next, we computed the mean values of subject-level occupancy rates (**Figure 8B)** and dwell times (**Figure 8C**) for each obtained cluster. Cluster 1, with the highest entropy across all networks, exhibits significantly greater occupancy rates for SZ patients than controls, while controls demonstrate significantly higher occupancy rates in clusters (2 and 4) which have low ICE values. Cluster 5 is characterized by high ICE values across all networks along with high occupancy for both groups. Mean dwell time for high-entropy cluster 1 is higher in SZ patients whereas low-entropy cluster 2 has higher mean dwell time for HCs. Clusters 3, 4, and 5 exhibit the same mean dwell time for both SZs and HCs. Despite strong effect of the diagnosis on mean dwell time in clusters 1 and 2 these results are not statistically significant after FDR correction (**Table S2**).

Also we calculated average ICE values across windows and individuals for two groups (**Figure 8D,E**) and their difference (**Figure 8F**). The clusters 2 and 4, which exhibit low entropy, demonstrate more distinct patterns of ICE in both SZ and control groups across various functional brain networks. The group difference (SZ–HC) in average DICE values for each cluster is shown in **Figure 8J**. Table with group difference p-values (**Table S3**) corresponding to each ICN and each cluster is presented in ‘Supplementary Materials’ section. Although most ICNs in low-entropy clusters 2 and 4 exhibit significant group differences in average ICE (**Figure 8F**), high-entropy clusters 1, 3, and 5 demonstrate statistically stronger results. This phenomenon is explained by the fact that the standard deviation for ICE in most ICNs is much higher in clusters 2 and 4 than in clusters 1, 3, and 5 (**Figure S1A**,**B**).

Furthermore, Cluster 4, which has the lowest ICE, exhibits higher standard deviation of ICE for healthy controls compared to schizophrenia patients for the majority of ICNs (**Figure S1C**).

Next, we examined the distributions of dynamic ICE values for each cluster (**Figure 9**). The histograms validate that the patterns in the centroids are highly characteristic of the cluster elements. Histograms for less-occupied clusters 2 and 4 exhibit a bimodal distribution and broader spread compared to the high-occupancy, high-entropy clusters 1 and 5, which are unimodal and narrowly distributed. Bimodality of distributions related to clusters 2 and 4 is in alignment with significantly higher STD of ICE for these clusters (**Figure S1A**,**B**). To compare SZ and HC dynamic ICE distributions we utilized the Kolmogorov-Smirnov (K-S) test. K-S rejected the null hypothesis at 5% significance level for all clusters. This means that SZ and HC distributions of dynamic ICE associated with given cluster are statistically different for all five clusters.

**Figure 9.**
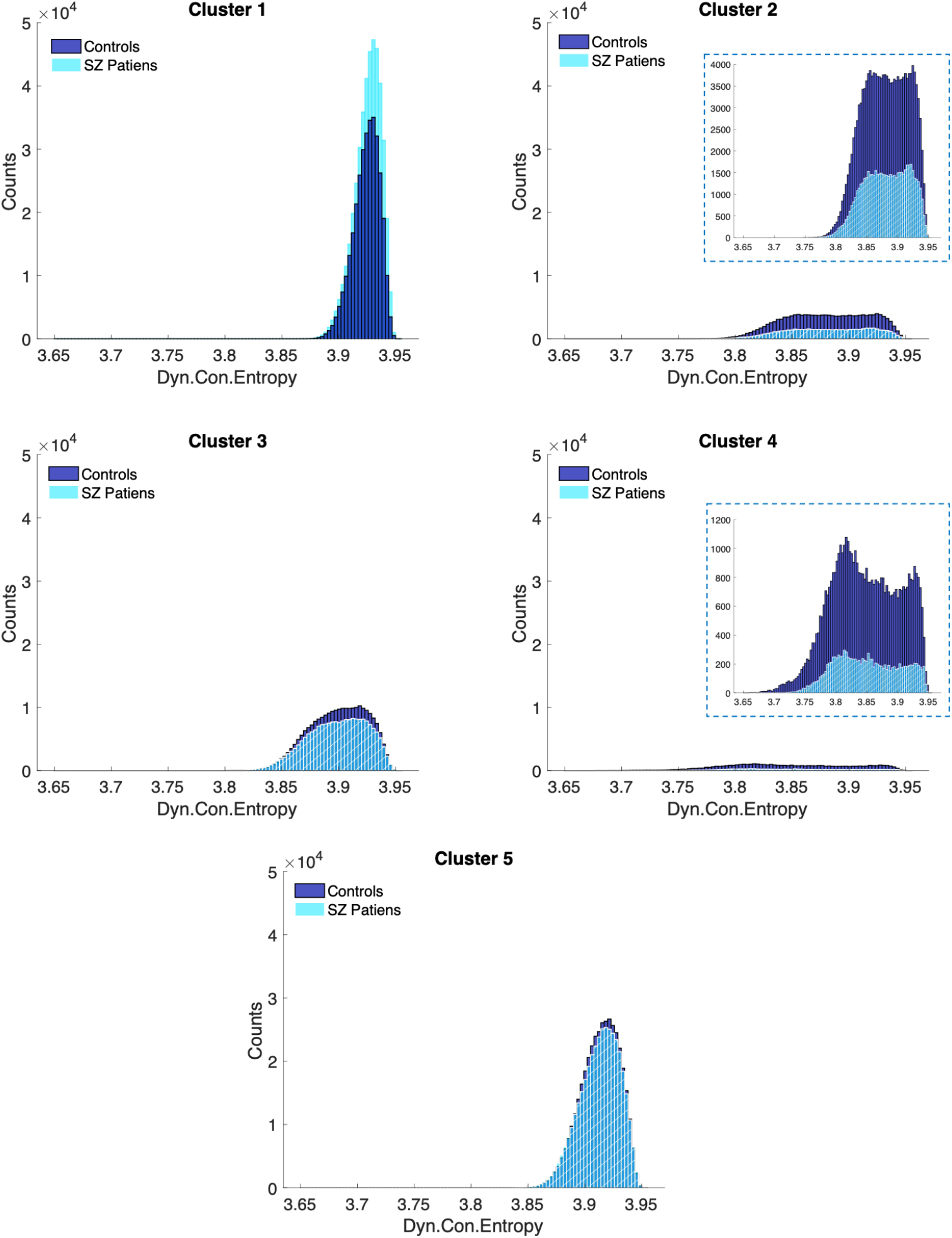
The histograms associated with less occupied low-entropy clusters 2 and 4 are bimodal and more broadly distributed, whereas histograms associated with high occupied high entropy clusters1 and 5 are unimodal and more narrowly distributed. For every histogram corresponding to a specific cluster of each group, we collapsed and aggregated the 53-length vectors present within the cluster, then showed how many of these individual elements from dynamic ICEs in this cluster are in each bin referenced on the x-axis.

## 4 Discussion

Our results demonstrated that with the proposed new measure – ICE, we were able to identify links to SZ among a range of functional brain domains: SC, AUD, VIS, SM, CC, DM and CB. All these domains showed higher mean ICE in SZ patients compared to HC. Higher ICE associated with individuals with SZ indicates that patients demonstrate higher randomness in distribution of time-varying connectivity strength across functional regions from each source network. This is consistent with, and extends, prior studies showing more randomness/disorganization in functional brain connectivity in SZ patients when compared to control group (He et al., 2012; Ramirez-Mahaluf et al., 2022) as well as with (Carhart-Harris et al., 2014) work that uses other entropy approaches not based on ICA and dFNC. According to (Carhart-Harris et al., 2014) brain entropy is suppressed during normal waking consciousness when brain operates just below criticality, however, at psychedelic state entropy is increased, particularly at hippocampus and anterior cingulate cortex, the networks that showed increased ICE in SZ patients in our study. It is known that SZ and psychedelics share similar effects on mental health, particularly in their neural activation patterns during hallucinations (Leptourgos et al., 2020).

In addition, our ICE metric revealed that SZ predominantly affected intrinsic functional brain networks associated with the SC, VIS, SM and CB, domains while approximately half of the AUD, CC and DM ICNs were impacted. (**Table1, Figure 7C**). Our findings align with prior research suggesting that individuals with SZ demonstrate pervasive alterations in perception and sensory processing, exhibit distorted thinking, and experience impaired cognitive functions (Kalkstein et al., 2010; Uhlhaas & Singer, 2010). Individuals with schizophrenia also exhibit disruptions in the mechanisms responsible for processing auditory (Dondé et al., 2019), visual (Adámek et al., 2022; Dondé et al., 2019), and somatosensory modalities and motor functions. Also, DMN has been widely observed to be abnormal in schizophrenia, and the mental processes associated with this network are pertinent to the disease (Hu et al., 2017; Zhou et al., 2015). Abnormal activity and functional connectivity in the DMN regions of SZ patients is also related to cognitive deficits and psychopathology related to the disease (Calhoun et al., 2011; Hu et al., 2017; Zhou et al., 2007).

Reduced ICE correlation between SC, AUD, VIS, SM and CB reflects hypoconnectivity between AUD, VIS and SM ICNs in SZ patients reported in study (Damaraju et al., 2014) and weaker connectivity between SC and CB ICNs in SZ patients in research (Soleimani et al., 2024) that uses same dataset as in our study. In addition, we revealed increased ICE correlation between CC (inferior parietal lobule) and DMN (all ICNs) in patients (**Figure 7C,D**). Particularly high ICE correlation was observed between Inferior parietal lobule (network 26 in CC domain) and Posterior cingulate cortex (network 49 in DM domain) in patients (**Figure 7D**).

Furthermore, we showed that SZ patients tend to have larger occupancy weights in clusters characterized by weak, low-scale functional entropy correlation, whereas the control group exhibits higher occupancy weights in clusters with strong, large-scale functional entropy correlation. These results are consistent with, but extend, FNC state difference between SZ patients and controls shown in (Damaraju et al., 2014) which demonstrated that clusters characterized by weak and low-scale functional connectivity have greater occupancy among SZ patients compared to HC, whereas clusters with strong and large-scale connectivity are predominantly occupied by HC rather than SZ patients.

Our research demonstrated that both static and dynamic mean ICE was higher in SZ patients than in healthy controls. Histograms of both static and dynamic ICE for SZ patients have a larger portion of the mass at the higher end of the distributions compared to controls. Nevertheless, dynamic ICE enabled us to find additional parameters that transiently discern SZ patients from controls. Thus, we showed that distributions of network connectivity strength across ICNs of patients are less variable in time maintaining relatively consistent levels of ICE compared to controls. In addition, dynamic ICE analysis enabled us to reveal that human brain can function in distinct states of ICE: states with uniformly high entropy in connectivity strength for all ICN (states 1 and 5) and states with relatively low and uneven entropy in connectivity strength across different brain networks (clusters 2 and 4) (**Figure 8A**). Individuals with SZ have larger occupancy rates for state 1 with highest ICE, whereas healthy controls have higher occupancy for low-entropy states 2 and 4 with more structured given network’s connectivity to all the other brain networks (**Figure 8B**). Moreover, states 1 and 5 with high entropy are largely occupied by both HCs and SZ when compared to low-entropy states 2 and 4. States with lower or mixed entropy are relatively rare, and significantly rarer in SZ patients. Thus, broadly speaking, dynamic ICE analysis reveals a prevailing tendency for the brain to be circulating through connectivity patterns with relatively high entropy levels, which aligns with or complexity in distribution of time-varying connectivity strength across functional brain networks. Thus, circulating through less organized/structured connectivity patterns as long as these fluctuations occasionally converge into more focused patterns appears healthy. In individuals with SZ, there seems to be some impediment preventing them from transiently achieving these more focused and structured connectivity patterns.

It is important to notice that many intrinsic functional brain networks exhibit the most noticeable group differences in states (1, 3, 5) where ICE is high for majority of ICNs (**Figure 8J**). Also, cluster 1 with highest DICE and highest occupancy and dwell time for SZ patients is an only cluster where all ICNs (except for Precuneus, ICN of DMN) have higher DICE for SZ patients than HCs. Particularly SC, SM, VIS and CB brain domains have significantly larger DICE in SZ patients compared to healthy controls. This tells us that SZ patients’ brain circulates mostly through more chaotic/less organized functional connectivity patterns. It is also crucial to observe inability of SZs to achieve states (cluster 2 and 4) in which the SC (particularly Subthalamus/Hypothalamus and Thalamus), VIS (particularly Middle temporal gyrus), and cerebellar networks specifically are not concentrating their connectivity in specific brain regions.

It is interesting to observe that both SZs and HCs exhibit the highest mean dwell time for high-entropy state 1 compared to other states. Thus, both SZs and HCs spend the majority of their time in high-entropy state 1, with patients having a higher mean dwell time. While SZs have a higher mean dwell time for state 1, and HC spend more time in state 2, the effect of diagnosis is strong but not statistically significant after FDR correction. The differences between HC and SZ in mean dwell time appear to be less significant than mean occupancy rates, suggesting that the rate at which the groups change states is more similar between HC and SZ than which states they change to.

Despite offering valuable insights into time-varying heterogeneity of brain network’s connectivity at healthy and disease state using a novel ICE approach the presented study has at least two limitations. First, the applicability of the findings may be limited by the specific dataset used, which included 311 participants, comprising 151 schizophrenia patients and 160 age and gender-matched healthy controls. Enlarging the sample size and introducing more diversity could offer a broader representation of the population and could improve the reliability of the findings. Second, this study does not account for common confounding factors such as the use of antipsychotics and other psychotropic medications, current smoking, and prior history of substance use. Further research is needed to consider these varying confounding factors. Also, it would be interesting to explore the relationships between ICE findings and illness characteristics, such as positive and negative symptoms, various cognitive deficits, and the duration of illness.

## 5 Conclusion

The proposed inter-network connectivity entropy (ICE) measure together with functional brain connectivity analyses appear to be simple and reliable way to summarize time-varying FNC data and investigate group effects for potential clinical application. In addition to the advantages of the time-varying whole-brain FNC approach—such as robustness, reproducibility, and freedom from constraints related to the selection of specific seeds or regions of interest—our approach provides a new level of understanding of both physiological and pathophysiological brain states. Firstly, both static and dynamic ICE measures showed that schizophrenia patients exhibit greater randomness/disorganization in the distribution of connectivity strength across various intrinsic connectivity networks spanning a wide range of functional brain domains, including subcortical, auditory, visual, sensorimotor, cognitive control, default mode, and cerebellar regions when compared to control group. Secondly, in general, the brains of schizophrenia patients are characterized by weak, low-scale functional entropy correlation across various functional brain regions, while healthy brains tend to show strong, large-scale functional entropy correlation. The dynamic ICE measure complements and extends our findings, revealing that, firstly, the healthy brain primarily navigates through complex, less focused connectivity patterns, with occasional transitions into more organized configurations of a given network’s connectivity to all other brain networks. However, schizophrenia patients’ brains circulate through more disorganized connectivity patterns compared to healthy controls and fail to achieve more focused functional connectivity patterns, especially evident in ICNs associated with subcortical (particularly subthalamus/hypothalamus and thalamus), visual (particularly middle temporal gyrus), and cerebellar brain domains which do not concentrate their connectivity in specific brain regions in individuals with schizophrenia. Secondly, ICE in schizophrenia patients shows significantly less variability over time compared to controls, suggesting lower temporal dynamics in functional connectivity strength distribution in patients. These insights highlight the potential applications of our methodology beyond schizophrenia. Our ICE measure can serve as the basis for a pipeline designed to classify and compare the impact of various diseases on the brain or to study the healthy brain and behavior relationships.

## 7 Data availability statement

Due to IRB restrictions, the FBIRN data analyzed in this study cannot be shared without specific licenses. However, the dataset can be accessed upon request by contacting Dr. Theo G.M. van Erp at, who will facilitate the interaction with the IRB.

## 10 Supporting Information

Additional supporting information can be found online in the Supporting Information section at the end of this article.

