## Supporting Information for "Static and Dynamic Cross-Network Functional Connectivity Shows Elevated Entropy in Schizophrenia Patients"

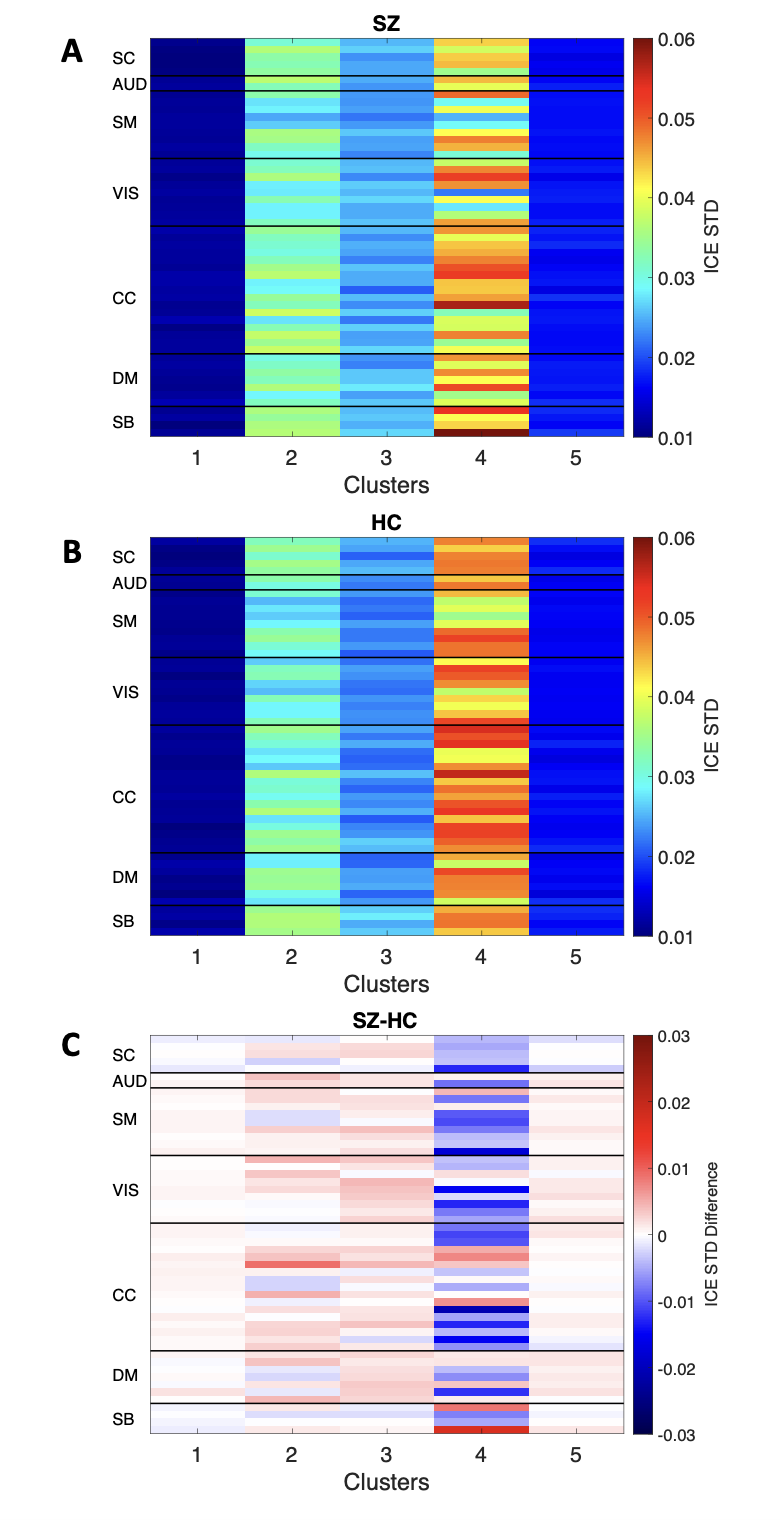

**Figure S1.** Low-entropy clusters 2 and 4 exhibit much higher standard deviations for ICE in both SZ (**A**) and HC (**B**), when compared to high-entropy clusters 1, 3, and 5. (**C**) Group differences in STD for DICE across ICNs and DICE clusters.

**Table S1**. Regression coefficients and p-values for diagnosis for both static and dynamic ICE associated with intrinsic connectivity networks. All significant results that have survived after FDR correction are displayed in bold.

| # | **Functional networks** | **SICE** | | **DICE** | |
| --- | --- | --- | --- | --- | --- |
|  |  | **coefficients** | **p-values** | **coefficients** | **p-values** |
|  | **Subcortical (SC)** |  | | | |
| 1 | Caudate | 0.0114 | **3.96x10^-8^** | 0.0143 | **1.26x10^-8^** |
| 2 | Subthalamus/hypothalamus | 0.0097 | **6.77x10^-7^** | 0.0135 | **1.80x10^-8^** |
| 3 | Putamen | 0.0026 | 3.94x10^-2^ | 0.0043 | **9.78x10^-3^** |
| 4 | Caudate | 0.0078 | **4.74x10^-5^** | 0.0101 | **1.69x10^-5^** |
| 5 | Thalamus | 0.0138 | **4.22x10^-9^** | 0.0177 | **2.55x10^-10^** |
|  | **Auditory (AUD)** |  | | | |
| 6 | Superior temporal gyrus | 0.0076 | **7.00x10^-5^** | 0.0087 | **1.26x10^-4^** |
| 7 | Middle temporal gyrus | 0.0005 | 7.02x10^-1^ | 0.0013 | 4.10x10^-1^ |
|  | **Sensorimotor (SM)** |  | | | |
| 8 | Postcentral gyrus | 0.0085 | **7.76x10^-5^** | 0.0099 | **6.95x10^-5^** |
| 9 | Left postcentral gyrus | 0.0090 | **4.67x10^-5^** | 0.0105 | **4.15x10^-5^** |
| 10 | Paracentral lobule | 0.0086 | **8.16x10^-6^** | 0.0103 | **4.90x10^-6^** |
| 11 | Right postcentral gyrus | 0.0096 | **3.06x10^-5^** | 0.0107 | **5.20x10^-5^** |
| 12 | Superior parietal lobule | 0.0075 | **1.08x10^-4^** | 0.0093 | **6.06x10^-5^** |
| 13 | Paracentral lobule | 0.0038 | **2.13x10^-2^** | 0.0051 | **8.36x10^-3^** |
| 14 | Precentral gyrus | 0.0038 | **2.13x10^-2^** | 0.0048 | **1.16x10^-2^** |
| 15 | Superior parietal lobule | 0.0034 | **8.99x10^-3^** | 0.0049 | **1.94x10^-3^** |
| 16 | Postcentral gyrus | 0.0016 | 1.91x10^-1^ | 0.0035 | **2.37x10^-2^** |
|  | **Visual (VIS)** |  | | | |
| 17 | Calcarine gyrus | 0.0108 | **2.61x10^-6^** | 0.0127 | **2.28x10^-6^** |
| 18 | Middle occipital gyrus | 0.0048 | **3.51x10^-3^** | 0.0067 | **1.32x10^-3^** |
| 19 | Middle temporal gyrus | 0.0088 | **1.66x10^-6^** | 0.0115 | **1.29x10^-7^** |
| 20 | Cuneus | 0.0101 | **5.07x10^-6^** | 0.0115 | **8.83x10^-6^** |
| 21 | Right middle occipital gyrus | 0.0089 | **8.08x10^-5^** | 0.0110 | **3.50x10^-5^** |
| 22 | Fusiform gyrus | 0.0037 | 5.15x10^-2^ | 0.0060 | **7.17x10^-3^** |
| 23 | Inferior occipital gyrus | 0.0091 | **4.58x10^-5^** | 0.0113 | **1.98x10^-5^** |
| 24 | Lingual gyrus | 0.0070 | **1.05x10^-3^** | 0.0087 | **5.78x10^-4^** |
| 25 | Middle temporal gyrus | 0.0036 | **1.77x10^-2^** | 0.0057 | **2.05x10^-3^** |
|  | **Cognitive Control (CC)** |  | | | |
| 26 | Inferior parietal lobule | 0.0000 | 9.78x10^-1^ | 0.0012 | 4.94x10^-1^ |
| 27 | Insula | 0.0040 | **5.41x10^-4^** | 0.0059 | **1.52x10^-4^** |
| 28 | Superior medial frontal gyrus | 0.0003 | 8.13x10^-1^ | 0.0021 | 1.76x10^-1^ |
| 29 | Inferior frontal gyrus | 0.0003 | 7.19x10^-1^ | 0.0017 | 1.76x10^-1^ |
| 30 | Right inferior frontal gyrus | 0.0012 | 2.06x10^-1^ | 0.0027 | **3.79x10^-2^** |
| 31 | Middle frontal gyrus | 0.0008 | 4.35x10^-1^ | 0.0025 | 6.94x10^-2^ |
| 32 | Inferior parietal lobule | 0.0036 | **2.74x10^-2^** | 0.0060 | **3.14x10^-3^** |
| 33 | Left inferior parietal lobule | 0.0013 | 2.73x10^-1^ | 0.0030 | 4.38x10^-2^ |
| 34 | Supplementary motor area | 0.0042 | **5.08x10^-5^** | 0.0053 | **8.97x10^-5^** |
| 35 | Superior frontal gyrus | -0.0017 | 1.42x10^-1^ | -0.0001 | 9.72x10^-1^ |
| 36 | Middle frontal gyrus | 0.0029 | **1.10x10^-2^** | 0.0054 | **4.97x10^-4^** |
| 37 | Hippocampus | 0.0041 | **8.20x10^-4^** | 0.0066 | **6.77x10^-5^** |
| 38 | Left inferior parietal lobule | -0.0003 | 7.34x10^-1^ | 0.0013 | 3.10x10^-1^ |
| 39 | Middle cingulate cortex | 0.0029 | 4.42x10^-2^ | 0.0046 | **1.07x10^-2^** |
| 40 | Inferior frontal gyrus | 0.0052 | **1.27x10^-3^** | 0.0076 | **1.74x10^-4^** |
| 41 | Middle frontal gyrus | 0.0066 | **8.56x10^-4^** | 0.0086 | **2.80x10^-4^** |
| 42 | Hippocampus | 0.0047 | **3.82x10^-4^** | 0.0077 | **8.44x10^-6^** |
|  | **Default Mode (DMN)** |  | | | |
| 43 | Precuneus | -0.0007 | 5.13x10^-1^ | 0.0006 | 6.91x10^-1^ |
| 44 | Precuneus | 0.0007 | 4.18x10^-1^ | 0.0015 | 2.33x10^-1^ |
| 45 | Anterior cingulate cortex | 0.0038 | **9.98x10^-3^** | 0.0062 | **8.95x10^-4^** |
| 46 | Posterior cingulate cortex | 0.0028 | 7.85x10^-2^ | 0.0044 | **2.09x10^-2^** |
| 47 | Anterior cingulate cortex | 0.0053 | **4.63x10^-4^** | 0.0082 | **2.40x10^-5^** |
| 48 | Precuneus | 0.0006 | 5.57x10^-1^ | 0.0017 | 2.07x10^-1^ |
| 49 | Posterior cingulate cortex | -0.0027 | **1.64x10^-2^** | -0.0009 | 5.32x10^-1^ |
|  | **Cerebellum (CB)** |  | | | |
| 50 | Cerebellum 1 | 0.0115 | **9.35x10^-8^** | 0.0135 | **8.15x10^-8^** |
| 51 | Cerebellum 2 | 0.0088 | **2.35x10^-5^** | 0.0113 | **4.27x10^-6^** |
| 52 | Cerebellum 3 | 0.0071 | **2.12x10^-6^** | 0.0092 | **1.40x10^-6^** |
| 53 | Cerebellum 4 | 0.0110 | **6.77x10^-8^** | 0.0136 | **2.98x10^-8^** |

**Table S2.** Statistics for mean occupancy rates and mean dwell time associated with five states of DICE.

p-values that survived after FDR correction are shown in bold.

| **DICE Clusters** | | **Regression coefficients for diagnosis** | **p-values** |
| --- | --- | --- | --- |
| Mean Occupancy Rate | State 1 | 0.149 | **3.87x10^-5^** |
|  | State 2 | -0.072 | **6.61x10^-6^** |
|  | State 3 | -0.038 | 8.10x10^-2^ |
|  | State 4 | -0.028 | **3.31x10^-3^** |
|  | State 5 | -0.011 | 6.31x10^-1^ |
| Mean Dwell Time | State 1 | 6.282 | 2.55x10^-2^ |
|  | State 2 | -2.412 | 4.02x10^-2^ |
|  | State 3 | -0.266 | 7.73x10^-1^ |
|  | State 4 | 0.107 | 9.69x10^-1^ |
|  | State 5 | -0.054 | 9.46x10^-1^ |

**Table S3.** Statistics for DICE associated with functionally relevant ICNs across different clusters. The results were obtained via a two-sample t-test followed by FDR correction. P-values corresponding to statistically significant differences in DICE between SZ patients and healthy controls, which have been FDR corrected, are displayed in bold.

| # | **Functional networks** | **DICE p-values** | | | | |
| --- | --- | --- | --- | --- | --- | --- |
|  |  | **State 1** | **State 2** | **State 3** | **State 4** | **State 5** |
|  | **Subcortical (SC)** |  | | | | |
| 1 | Caudate | **8.75x10^-118^** | **5.43x10^-12^** | **3.85x10^-29^** | **5.18x10^-17^** | **1.22x10^-58^** |
| 2 | Subthalamus/hypothalamus | **2.70x10^-16^** | **4.65x10^-53^** | **5.67x10^-06^** | **1.16x10^-32^** | **5.58x10^-21^** |
| 3 | Putamen | **2.46x10^-08^** | **4.20x10^-12^** | **2.65x10^-17^** | 3.33x10^-01^ | **5.15x10^-12^** |
| 4 | Caudate | **4.88x10^-14^** | **2.34x10^-09^** | **3.85x10^-03^** | **2.99x10^-09^** | **2.48x10^-03^** |
| 5 | Thalamus | **4.89x10^-81^** | **2.22x10^-70^** | **1.94x10^-104^** | **7.41x10^-22^** | **9.52x10^-170^** |
|  | **Auditory (AUD)** |  | | | |  |
| 6 | Superior temporal gyrus | **6.24x10^-13^** | **2.89x10^-02^** | **5.45x10^-12^** | **6.86x10^-04^** | **3.56x10^-11^** |
| 7 | Middle temporal gyrus | **8.32x10^-04^** | **1.31x10^-07^** | **9.56x10^-09^** | **9.57x10^-07^** | **2.56x10^-26^** |
|  | **Sensorimotor (SM)** |  | | | |  |
| 8 | Postcentral gyrus | **4.63x10^-11^** | **1.02x10^-10^** | **3.05x10^-06^** | **2.80x10^-05^** | **2.61x10^-11^** |
| 9 | Left postcentral gyrus | **6.14x10^-11^** | **2.59x10^-03^** | 7.74x10^-01^ | 7.78x10^-01^ | 5.73x10^-02^ |
| 10 | Paracentral lobule | **2.20x10^-26^** | **1.28x10^-04^** | **3.22x10^-04^** | 5.69x10^-01^ | 3.63x10^-01^ |
| 11 | Right postcentral gyrus | **4.42x10^-07^** | 8.48x10^-02^ | **5.55x10^-03^** | **8.33x10^-03^** | **5.43x10^-03^** |
| 12 | Superior parietal lobule | **7.07x10^-27^** | **1.07x10^-04^** | 5.87x10^-02^ | **9.59x10^-03^** | 7.42x10^-02^ |
| 13 | Paracentral lobule | **1.80x10^-22^** | **2.75x10^-04^** | **4.66x10^-33^** | 9.26x10^-02^ | **6.05x10^-13^** |
| 14 | Precentral gyrus | **1.63x10^-23^** | 3.86x10^-01^ | **1.08x10^-38^** | 2.96x10^-01^ | **2.39x10^-23^** |
| 15 | Superior parietal lobule | 5.44x10^-02^ | 5.14x10^-01^ | **1.60x10^-08^** | **8.44x10^-11^** | **3.17x10^-07^** |
| 16 | Postcentral gyrus | **2.05x10^-04^** | 3.54x10^-02^ | **8.69x10^-04^** | 9.11x10^-01^ | **2.00x10^-02^** |
|  | **Visual (VIS)** |  | | | |  |
| 17 | Calcarine gyrus | **2.08x10^-59^** | **2.65x10^-07^** | **1.71x10^-18^** | **2.57x10^-02^** | **7.30x10^-11^** |
| 18 | Middle occipital gyrus | **2.32x10^-04^** | 5.89x10^-01^ | **2.25x10^-08^** | **1.18x10^-03^** | **9.85x10^-08^** |
| 19 | Middle temporal gyrus | **5.30x10^-39^** | **2.60x10^-21^** | **2.12x10^-28^** | **8.88x10^-22^** | **6.00x10^-24^** |
| 20 | Cuneus | **4.93x10^-31^** | 1.93x10^-01^ | **5.92x10^-08^** | 4.35x10^-01^ | **8.21x10^-23^** |
| 21 | Right middle occipital gyrus | **3.99x10^-28^** | 6.27x10^-01^ | 2.67x10^-01^ | 7.56x10^-01^ | **3.96x10^-06^** |
| 22 | Fusiform gyrus | **1.22x10^-02^** | 1.57x10^-01^ | **2.97x10^-25^** | **4.87x10^-02^** | **1.22x10^-31^** |
| 23 | Inferior occipital gyrus | **5.55x10^-55^** | 3.48x10^-01^ | 1.66x10^-01^ | 1.06x10^-01^ | **2.32x10^-08^** |
| 24 | Lingual gyrus | **3.51x10^-06^** | **2.12x10^-02^** | **3.13x10^-06^** | 6.07x10^-02^ | **8.03x10^-05^** |
| 25 | Middle temporal gyrus | 1.42x10^-01^ | **1.07x10^-04^** | **2.09x10^-08^** | **3.77x10^-13^** | **4.41x10^-30^** |
|  | **Cognitive Control (CC)** |  | | | |  |
| 26 | Inferior parietal lobule | 5.77x10^-01^ | **1.42x10^-08^** | **5.85x10^-36^** | **2.89x10^-05^** | **1.86x10^-45^** |
| 27 | Insula | **5.56x10^-14^** | **3.34x10^-05^** | **6.68x10^-03^** | **4.69x10^-05^** | **3.36x10^-04^** |
| 28 | Superior medial frontal gyrus | 1.12x10^-01^ | 3.48x10^-02^ | **4.06x10^-18^** | **2.14x10^-02^** | **1.54x10^-14^** |
| 29 | Inferior frontal gyrus | **1.24x10^-19^** | **5.43x10^-05^** | **3.72x10^-24^** | **4.42x10^-13^** | 5.83x10^-01^ |
| 30 | Right inferior frontal gyrus | **2.44x10^-12^** | **4.09x10^-06^** | **2.12x10^-08^** | **1.14x10^-03^** | **1.64x10^-02^** |
| 31 | Middle frontal gyrus | **6.87x10^-08^** | **1.28x10^-15^** | **1.45x10^-28^** | 1.28x10^-01^ | **9.45x10^-03^** |
| 32 | Inferior parietal lobule | **4.69x10^-14^** | **1.71x10^-03^** | 7.67x10^-01^ | **4.17x10^-05^** | 5.47x10^-02^ |
| 33 | Left inferior parietal lobule | **8.87x10^-06^** | **3.79x10^-07^** | **3.82x10^-08^** | 3.33x10^-01^ | **1.76x10^-02^** |
| 34 | Supplementary motor area | **7.89x10^-11^** | **3.70x10^-03^** | **1.28x10^-07^** | **1.92x10^-10^** | **9.09x10^-10^** |
| 35 | Superior frontal gyrus | 2.55x10^-01^ | **9.16x10^-21^** | **4.55x10^-21^** | **1.31x10^-06^** | **2.47x10^-19^** |
| 36 | Middle frontal gyrus | **3.49x10^-03^** | 6.05x10^-02^ | **1.02x10^-02^** | 6.99x10^-02^ | **3.69x10^-06^** |
| 37 | Hippocampus | 9.12x10^-01^ | 4.62x10^-01^ | **9.86x10^-10^** | **6.05x10^-05^** | 8.52x10^-01^ |
| 38 | Left inferior parietal lobule | 4.18x10^-01^ | 5.35x10^-01^ | **1.25x10^-24^** | 5.26x10^-02^ | **1.77x10^-14^** |
| 39 | Middle cingulate cortex | **6.28x10^-08^** | **7.87x10^-12^** | **4.78x10^-09^** | **2.80x10^-04^** | **3.71x10^-18^** |
| 40 | Inferior frontal gyrus | 9.48x10^-02^ | 7.31x10^-02^ | **2.16x10^-02^** | **2.40x10^-03^** | 8.43x10^-01^ |
| 41 | Middle frontal gyrus | **2.17x10^-03^** | 6.78x10^-01^ | **1.65x10^-11^** | **1.70x10^-03^** | 7.98x10^-01^ |
| 42 | Hippocampus | **6.64x10^-10^** | **7.01x10^-05^** | **1.43x10^-03^** | **2.17x10^-04^** | **1.46x10^-08^** |
|  | **Default Mode (DMN)** |  | | | |  |
| 43 | Precuneus | 3.03x10^-01^ | **1.32x10^-02^** | **3.04x10^-22^** | **3.88x10^-10^** | **9.02x10^-39^** |
| 44 | Precuneus | **2.53x10^-09^** | **7.24x10^-07^** | **1.77x10^-15^** | 5.77x10^-01^ | **4.01x10^-12^** |
| 45 | Anterior cingulate cortex | **2.39x10^-19^** | 4.89x10^-01^ | 7.62x10^-01^ | 4.38x10^-01^ | **2.31x10^-10^** |
| 46 | Posterior cingulate cortex | **8.49x10^-03^** | **1.08x10^-02^** | **1.60x10^-03^** | **2.65x10^-02^** | **3.63x10^-16^** |
| 47 | Anterior cingulate cortex | **7.62x10^-21^** | 4.09x10^-02^ | **1.54x10^-11^** | **8.27x10^-07^** | 2.64x10^-01^ |
| 48 | Precuneus | **1.31x10^-11^** | 3.62x10^-02^ | **2.90x10^-38^** | 4.64x10^-01^ | **1.35x10^-44^** |
| 49 | Posterior cingulate cortex | **2.41x10^-05^** | **7.63x10^-10^** | **1.46x10^-22^** | **1.20x10^-11^** | **3.86x10^-18^** |
|  | **Cerebellum (CB)** |  | | | |  |
| 50 | Cerebellum 1 | **3.28x10^-23^** | **3.86x10^-27^** | **1.95x10^-22^** | **6.03x10^-07^** | **1.30x10^-23^** |
| 51 | Cerebellum 2 | **2.19x10^-36^** | **4.65x10^-05^** | **1.92x10^-14^** | **1.04x10^-06^** | **7.00x10^-15^** |
| 52 | Cerebellum 3 | **8.08x10^-26^** | **2.48x10^-08^** | **6.90x10^-03^** | **3.02x10^-22^** | **1.00x10^-06^** |
| 53 | Cerebellum 4 | **2.20x10^-39^** | **1.65x10^-13^** | **6.49x10^-20^** | **4.68x10^-11^** | **6.32x10^-41^** |
